# Two-Stage Multivariate Mendelian Randomization on Multiple Outcomes with Mixed Distributions

**DOI:** 10.1101/2022.05.29.493904

**Authors:** Yangqing Deng, Dongsheng Tu, Chris J O’Callaghan, Geoffrey Liu, Wei Xu

## Abstract

In clinical research, it is of importance to study whether certain clinical factors or exposures have causal effects on clinical and patient reported outcomes like toxicities, quality of life, and self-reported symptoms, which can help improve patient care. Usually, such outcomes are recorded as multiple variables with different distributions. Mendelian randomization is a commonly used technique for causal inference with the help of genetic instrumental variables to deal with observed and unobserved confounders. Nevertheless, the current methodology of Mendelian randomization on multiple outcomes only focuses on one outcome at a time, meaning that it does not consider the correlation structure of multiple outcomes, which may lead to loss of statistical power. In situations with multiple outcomes of interest, especially when there are mixed correlated outcomes with multiple distributions, it is much more desirable to jointly analyze them with a multivariate approach. Some multivariate methods have been proposed to model mixed outcomes, however, they do not incorporate instrumental variables and cannot handle unmeasured confounders. To overcome the above challenges, we propose a two-stage multivariate Mendelian Randomization Method (MRMO), that can perform multivariate analysis on mixed outcomes using instrumental variables. We demonstrate that our proposed MRMO algorithm can gain power over the existing univariate method through simulation studies and a clinical application on a randomized Phase III clinical trial study on colorectal cancer patients.

## 1. Introduction

With the increasing collection of data on various clinical and patient reported outcomes (e.g., toxicities and quality of life measures) in clinical and observational studies, there is a growing interest in detecting the causal relationship between an exposure variable (X) and multiple outcomes of interest (Y) (Ringash and others, 2013; Friese and others, 2017; Kolotkin and Andersen, 2017; Patrinely and others, 2020; Martinsen and others, 2021). For example, some baseline clinical characteristics may affect a patient’s risk of experiencing certain toxicities after receiving a treatment. Finding and studying such causal effects may help clinicians predict the risk of adverse events and make better treatment decisions (He and others, 2014; Gowen and others, 2018). This can be illustrated through the analysis of data from CO.20 trial conducted by the Canadian Cancer Trials Group, which was a phase III randomized, placebo-controlled study aiming to examine the effect of the addition of brivanib (BRI) to cetuximab on colorectal cancer patients (Siu and others, 2013; Ringash and others, 2013; Shepshelovich and others, 2018). The original analysis of the original showed that compared to the cetuximab plus placebo treatment, cetuximab plus BRI was associated with increased toxicities and did not significantly improve the overall survival of patients (Siu and others, 2013). It is of interest in whether certain baseline variables have any effect on the a few selected toxicities a patient may experience when going through the cetuximab plus placebo treatment, with the mixed-type measurements for these toxicities. There are two major challenges, however, in these analyses. One challenge is that, although patients may be randomized by treatment, patients may not be balanced by the baseline variable X of interest and, therefore, directly modelling the relationship between X and Y without considering the confounders, measured or unmeasured, may lead to misleading results (Angrist and others, 1996). The other challenge is that sometimes the multiple outcomes of interest may be correlated, and mixed with different types of distributions (e.g., binary and continuous) (Olkin and Tate, 1961). Studying each outcome separately is a commonly used straightforward approach but will not utilize the correlation information, while modelling mixed outcomes together may use more information but require more complex models (Sammel and others, 1999; Teixeira-Pinto and Normand, 2009).

From a clinical standpoint, there is a major analytical gap in our ability to study the influence of a causal effect on multiple outcomes simultaneously, particularly in the context of toxicities. In the areas of rheumatology, immunology/transplantation, and oncology, it is common for patients and physicians to accept multiple moderate to severe toxicities in their treatments, as the treatments are considered life-saving. Most of the drugs used in these settings have widely variable toxicities from person to person. Understanding etiological factors that lead to significant toxicities in some patients but not others is fundamental to precision medicine. Innovative methods for analyzing individual etiological factors on multi-dimensional toxicities, particular those that are graded differently (e.g. ordinal symptoms vs continuous laboratory-based toxicities) are greatly needed.

Facing the challenge of confounders, people have been applying the Mendelian randomization (MR) approach, which uses genetic variants as instrumental variables (Burgess and others, 2015). Under certain instrumental variable assumptions, MR methods can efficiently estimate and make inference on the causal effect of the exposure on a single outcome. Different MR methods have been proposed to address a variety of problems, such as how to handle invalid instruments and how to incorporate multiple exposures (Bowden and others, 2015; Bowden and others, 2016; Burgess and Thompson, 2015; Burgess and others, 2018; Sanderson and others, 2018; Sanderson, 2020). Nevertheless, we are not aware of any existing MR techniques that allow the researchers to jointly model mixed outcome variables. When there are multiple outcomes to be examined, people usually conduct univariate analysis, which analyzes each outcome separately. This may not be ideal for testing the overall hypothesis, whether the exposure has any effect on any outcome, since the correlation structure between different outcomes is not considered, and Bonferroni correction is usually required, which is known to be conservative (Moran, 2003).

To address the above issues, it is desirable to develop a multivariate MR approach that can handle mixed outcomes. Different methods have been proposed to directly model mixed outcomes jointly (Cox, 1972; Cox and Wermuth, 1992; Olkin and Tate, 1961; Gueorguieva and Agresti, 2001; Zhang and others, 2018), while it is computationally challenging to implement most of them into the MR framework, especially when the full likelihood is considered. Bai and others (2020) proposed to use a composite-likelihood method with the Newton-Rapson algorithm to alleviate the computation issue (Atkinson, 1989). All these existing methods only model the relationship between X and Y without using instrumental variables, which means they cannot handle unmeasured confounders.

We develop a two-stage multivariate MR method for mixed outcomes, denoted by MRMO, that combines the two-stage MR approach with the composite-likelihood based mixed response model to assess the causal effects of the exposure on different outcomes. We propose to use the gradient descent algorithm when jointly modeling mixed outcomes, which has been shown to have a number of advantages over the commonly used Newton-Rapson algorithm (Press and others, 1988; Strutz 2010; Pascanu and others, 2014). In terms of testing the overall hypothesis, it is natural to consider the Wald test for multivariate models. Besides that, we propose to conduct the aSPU test for multiple outcomes (Pan and others, 2014; Kim and others, 2015), which can combine a class of powered sum tests to gain power in certain scenarios. Through extensive simulations and an application on a randomized Phase III clinical trial study (CO.20) on colorectal cancer patients (Siu and others, 2013; Ringash and others, 2013; Shepshelovich and others, 2018), we demonstrate that compared to the commonly used univariate MR analysis, our proposed multivariate approach is able to improve the power of testing the overall hypothesis while controlling type I errors, which can help clinicians find potential associations and generate new hypotheses.

## 2. Methods

### 2.1. Univariate Model for Mendelian Randomization with Multiple Outcomes

In this section, we illustrate the basic framework of traditional MR analysis, depicted by Figure 1(A). Suppose we are interested in studying the causal relationship between an exposure (X) and a single outcome (Y_1_). Directly modelling Y_1_ versus X may give us very problematic results, since there may be confounders (U) that are unmeasured or not included in the model. MR analysis incorporates some genetic variants G, typically SNPs (single-nucleotide polymorphisms), to serve as instrumental variables (IVs) for causal inference. Using IVs can help us avoid the confounding problem, if certain IV assumptions are met (i.e., IVs should be associated with X; IVs should not affect or be affected by U; IVs should not affect Y_1_ through any pathways other than X).

**Fig. 1.**
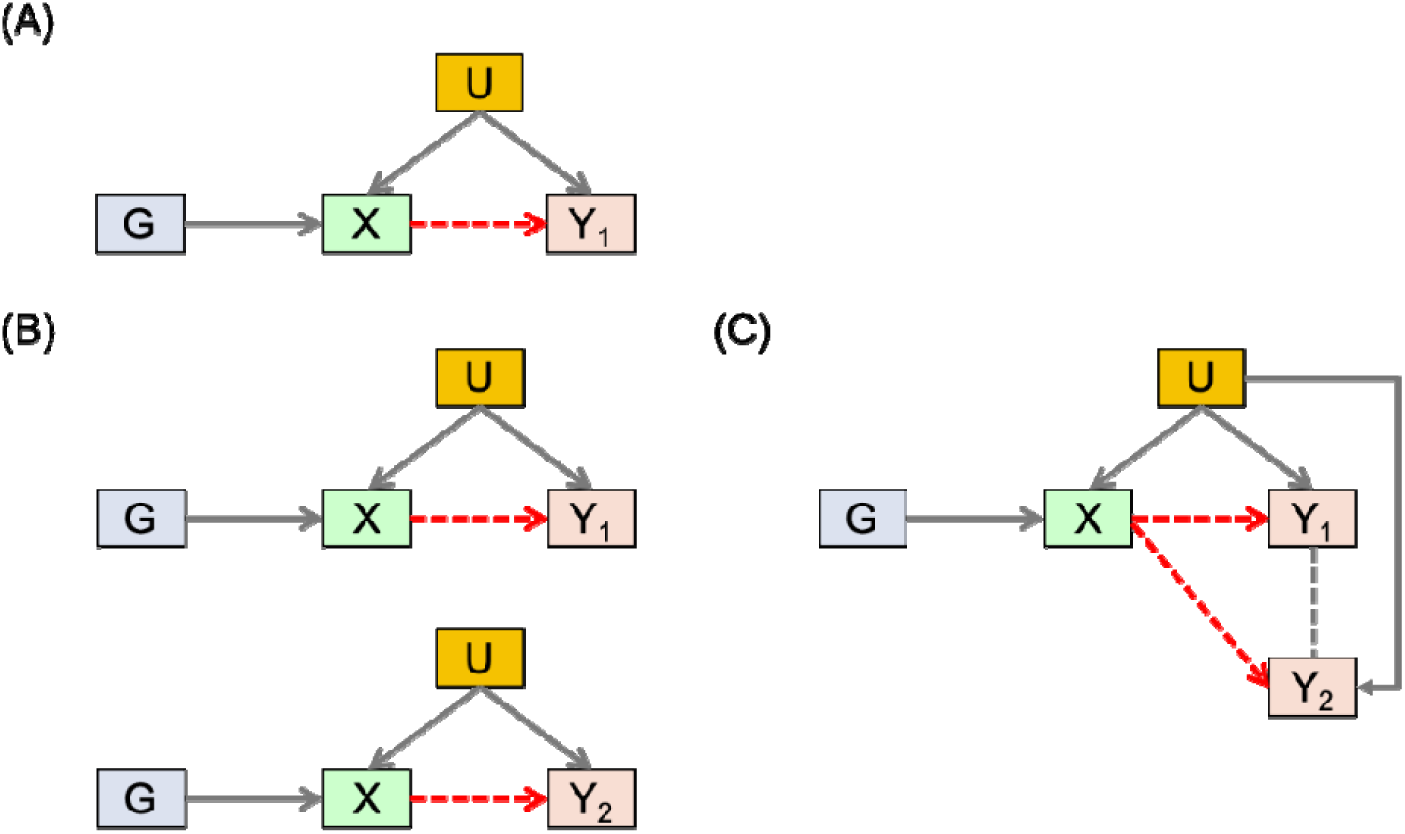
Basic framework. (A) MR analysis with a single outcome. (B) MR with multiple outcomes and univariate analysis. (C) MR with multiple outcomes and multivariate analysis.

When there are multiple outcomes of interest (e.g., different type of toxicities in cancer trials), people normally conduct several univariate analyses separately, which applies MR to one outcome at a time, as depicted by Figure 1(B) for two outcomes Y_1_ and Y_2_. However, looking at each outcome separately means ignoring the correlation information from the multiple outcomes, which may lead to loss of power, especially when we want to test the overall null hypothesis: the exposure does not affect any of the outcomes. Meanwhile, multivariate analysis, illustrated by Figure 1(C), models and analyzes multiple outcomes in a single model simultaneously. It may give us a better framework for testing the null hypothesis while accounting for the correlation structure of the different outcomes.

Before presenting our multivariate approach, we would like to give a brief overview of the two-stage univariate MR analysis using individual level data in a two-sample scenario, where we have two different datasets with different subjects for the exposure and outcomes. Suppose we look at *p* IVs and *q* outcomes, and the first sample set contains *n*^*^ subjects with 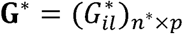 and 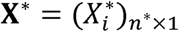, where 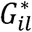 and 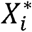 are the *l*th genotype value and the exposure value respectively for the *i*th subject in the first sample. The second sample set contains *n* subjects with **G** = (*G*_*il*_)_*n*×*q*_ and **Y** = (*Y*_*il*_)_*n*×*p*_, where *G*_*ij*_ and *Y*_*ij*_ are the *l*th genotype value and *j*th outcome value respectively for the *i*th subject in the second sample set. In the two-sample scenario, the *n*^*^ subjects and *n* subjects in the two sample sets do not overlap.

The first-stage regression model for the univariate MR is

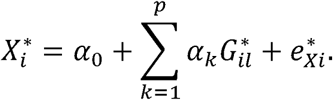

The first-stage coefficient estimates, denoted by 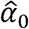 and 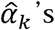, are used to predict the exposure value in the second sample, with 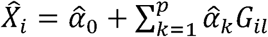. Then the second-stage regression models are

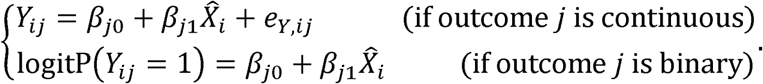

Note that for each outcome *j*, a single regression model is fitted separately, and the correlation structure between different outcomes is ignored. There are possible variations of the two-stage univariate MR analysis, including the adjusted two-stage method. Literature has shown that the two-stage method with residual inclusion, denoted by 2SRI, may be more suitable for binary outcomes than the standard two-stage method without residual inclusion (Terza and others, 2008). However, the standard method is more commonly used in practice and can provide a valid test of the association between the exposure and each outcome (Burgess and others, 2015). A study by Xue and Pan (2019) has also shown that the standard method may perform well and even better than 2SRI in certain scenarios, given that the effect sizes of the SNPs are usually small. For simplicity, in this article, we only focus on the standard method, which is more commonly used and can provide a valid test of the association between the exposure and each outcome (Burgess and others, 2015). The p-value of testing the exposure effect on a single outcome *j* is simply the p-value of the corresponding coefficient *β*_*j*1_. For binary outcomes, we also consider replacing logistic regression with probit regression, as it also allows us to test the association and is more comparable to the mixed response model we will describe, which uses the probit link.

### 2.2. Multivariate Mixed Response Model for Mendelian Randomization

In this section, we propose our two-stage multivariate MR method for mixed outcomes, which is expected to be more efficient than the univariate method when testing the overall null hypothesis (the exposure does not affect any of the outcomes). The first stage model is the same as what we have described in the univariate two-stage MR method, of which the main purpose is to use genetic information to predict the exposure for the second sample. In the second stage, instead of modeling each outcome separately, we propose to model different outcomes jointly using a multivariate mixed response model (Bai and others, 2020; Zhang and others, 2018). The relationship between each outcome and the fitted exposure can be modelled using a generalized linear model

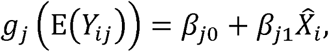

where the link function *g*_*j*_ can be chosen as the identity function for continuous outcomes, and the logistic or probit function for binary outcomes.

To model all outcomes simultaneously while incorporating their correlation structure, applying a conventional likelihood-based approach is possible but computationally very challenging. Using the pairwise composite likelihood approach as described in Bai and others (2020) may be a better option. For each pair of outcomes (*j,k*), the pairwise likelihood function can be written as

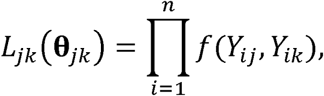

Where **θ**_*jk*_ represents the relevant parameters for *L*_*jk*_·*f*(*Y*_*ij*_,*Y*_*ik*_) is the pairwise likelihood for a single subject *i* only considering the joint distribution of outcomes *j,k*, which we will discuss shortly. Then the pairwise composite likelihood should be

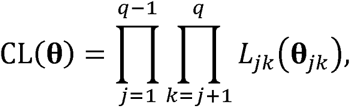

Where **θ** represents all of the parameters (Lindsay 1988; Cox & Reid 2004). The composite score function is

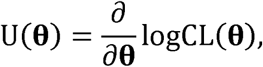

which means we can estimate the parameters by solving U (**θ**) = 0. Now we only need to specify *f*(*Y*_*ij*_,*Y*_*ik*_) properly according to the data types.

If a pair of outcomes (*j,k*) are both continuous, we assume they follow

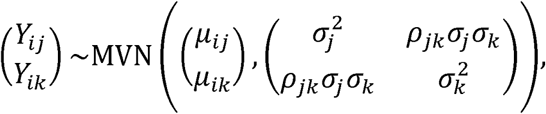

Where 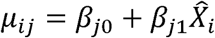 and 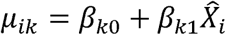, which means **θ**_*jk*_ consists of *β*_*j*0_, *β*_*j*1_, *β*_*k*0_, *β*_*k1*_, *σ*_*j*_, *σ*_*k*_ and *ρ*_*jk*_. Based on the density of a bivariate normal distribution, we have

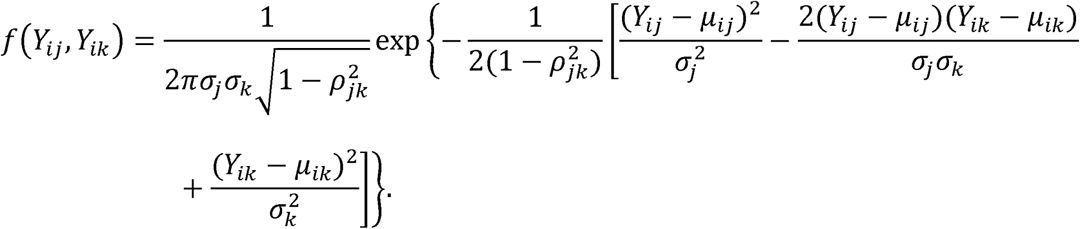

If a pair of outcomes (*j,k*) are both binary, following Bai and others (2020), we set up a pair of latent variables (*Z*_*ij*_,*Z*_*ik*_)and assume

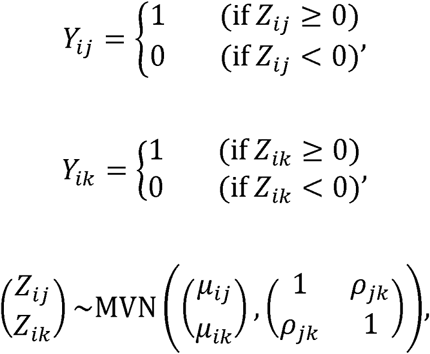

Where 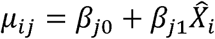 and 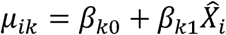. Note that modelling *Y*_*ij*_ this way is similar to using a probit regression model. Since (*Z*_*ij*_ − *µ*_*ij*_) follows a standard normal distribution, we know P(*Y*_*ij*_ = 1) = P(*Z*_*ij*_ ≥ 0)= P(*Z*_*ij*_ − *µ*_*ij*_ ≥ − *µ*_*ij*_) = Φ(*µ*_*ij*_) with Φ being the cumulative distribution function of the standard normal distribution. As a result, we have probitP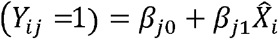. Let *s*_*ij*_ = 2*Z*_*ij*_ −1, *s*_*ik*_ = 2*Z*_*ik*_ −1. According to Bai and others (2020), we have

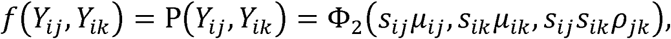

where Φ_2_ is the bivariate normal cumulative density function.

If one outcome *j* is binary, and another outcome *k* is continuous, we assume

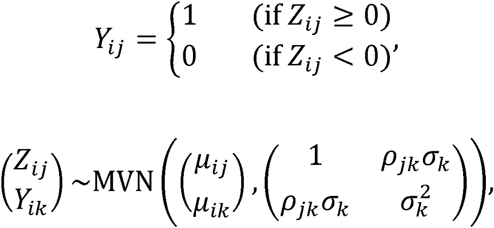

which gives us

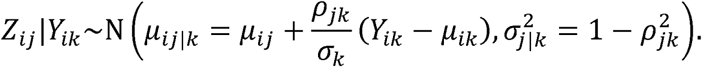

It can be derived that

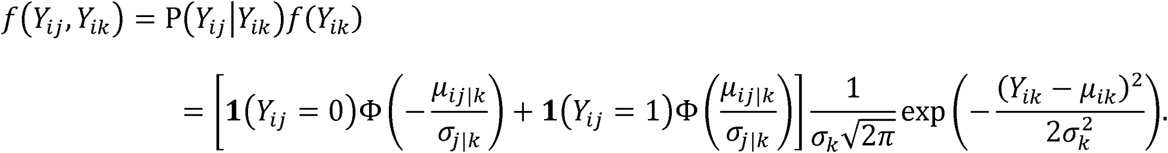

As a result, we can calculate *f* (*Y*_*ij*,_ *Y*_*ik*_) for any pair of outcomes given **θ**, and we can find the best **θ** by solving U(**θ**) = 0. To perform hypothesis testing, we also need to obtain the covariance matrix of the coefficient estimates 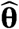. Based on the asymptotic theories for the composite likelihood function (Varin and others, 2011; Gao & Song 2010; Godambe 1960; Bai and others, 2020), the covariance matrix Cov 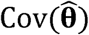 can be estimated as

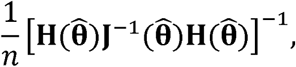

where

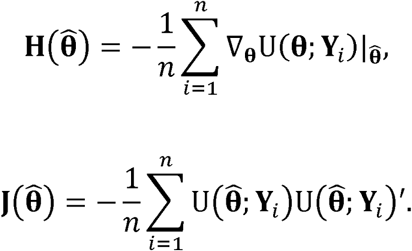

U (**θ**; **Y**_*i*_)denotes the score for subject *i*, which can be written as

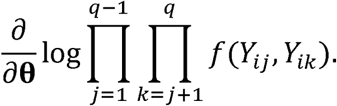

To estimate **θ**, we can use the traditional Newton–Raphson method, sometimes called Newton’s method, as suggested by Bai and others (2020) to solve U(**θ**) = 0. However, as explored by several researchers (Press and others, 1988; Strutz 2010; Pascanu and others, 2014), this algorithm relies heavily on the computation of the second derivative and may encounter problems like saddle points. If the initial estimates are far from the true parameters, Newton’s method may not work well. Hence, we propose to apply the gradient descent approach, which is more widely used in machine learning (Bottou, 2010; Amiri and Gunduz, 2019; Sun and others,2020). Aiming to find 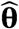 that minimizes logCL(**θ**), our algorithm is described below:

1. Choose the initial values of the parameters, denoted by **θ**^(0)^. A simple but effective option is to use the marginal model estimates based on the approach described in section 2.1.
2. Apply the gradient descent method to update the parameter estimates. For iteration *t* = 0,1,2, …, we have

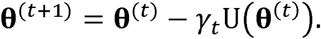
3. Calculate the new score function using the updated parameters.
4. Repeat the previous two steps until convergence. By default, the algorithm is stopped when ║**θ**^(*t*+1)^ − **θ**^(*t*)^ ║_1_ ≤ 1 /*n*.

Note that the choice of step size *γt* is crucial to the gradient descent algorithm. We recommend using the Barzilai–Borwein (BB) method, which is relatively straightforward and performs well in most scenarios (Barzilai and Borwein 1988; Fletcher, 2001). According to the BB method, the step size is chosen as

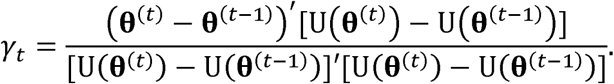

After obtaining 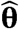, we can estimate the covariance matrix 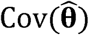 based on the previously described formulas. Then we can carry out the Wald test to test different null hypotheses, including whether the exposure affects a certain outcome *j* (H_0*j*_: *β*_*j*1_ = 0), and whether the exposure affects any outcome (H_0_: *β*_11_ = *β*_21_ = … = *β*_*q*1_ = 0).

### 2.3. Testing the Overall Hypothesis

In this section, we discuss more on the different choices for testing the overall hypothesis (H_0_: *β*_11_ = *β*_21_ = … = *β*_*q*1_ = 0), using univariate or multivariate methods. For the univariate analysis, the minP test with Bonferroni correction is usually applied. Suppose the p-values for the exposure effects on different outcomes are *p*_UVA,1_, *p*_UVA,2_, …, *p*_UVA,*q*_. The p-value for the overall test is (*qp*_UVA,1_, *qp*_UVA,2_, …, *qp*_UVA,*q*_,1) This method does not take into account the correlation between different outcomes, and it tends to be conservative.

For the multivariate analysis, once we have 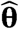 and 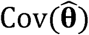, we can conduct the Wald test. From 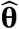 and 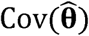, we can easily extract 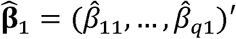 and 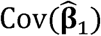. Then the test statistic is

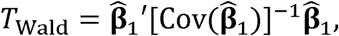

which approximately follows a chi-square distribution with *q* degrees of freedom under H_0_. Meanwhile, we propose to apply the aSPU test as an alternative method, which can combine a class of tests and may gain power when testing H_0_ (Pan and others, 2014; Kim and others, 2015). Denote 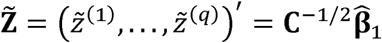, where **C** is a diagonal matrix with the same diagonal elements as 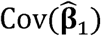. Under H_0_, 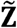 should asymptotically follow 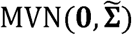, where 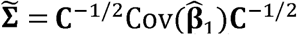. The overall null hypothesis can be examined by testing whether all of the Z-scores in 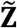 are 0. Define

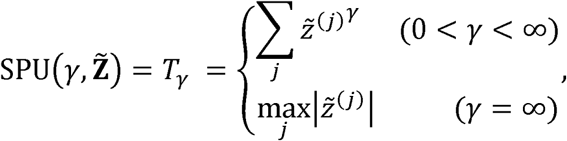

where each different choice of *γ* gives us a different test statistic by summing the powered Z-scores. Under the null, the Z-scores should have mean 0, and the sum of powered Z-scores should be relatively small. If the sum is too big, we should reject the null hypothesis. We consider a set of different *γ*’s, denoted by Γ = { *γ*_1_, *γ*_2_… *γ*_*r*_}, which is usually chosen as {1, 2, …, 8, ∞}. To obtain the p-value of 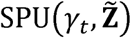, we sample 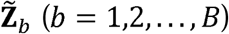 from the null distribution 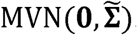, and then we have

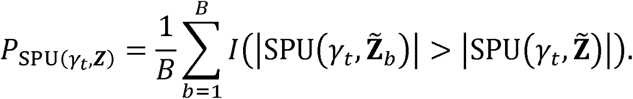

As a result, we can obtain a set of p-values 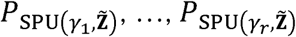, The aSPU test statistic is defined as 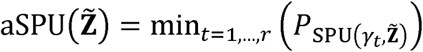, The general idea is that as long as one of the p-values is small enough, we should reject the null hypothesis, so we only need to examine whether 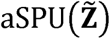 is too small. To obtain the p-value of 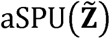, we need to use the empirical distribution of the aSPU test statistic under the null. For each 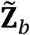, the aSPU test statistic can be calculated using

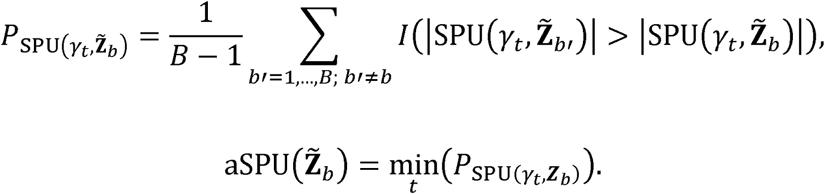

Hence, we can obtain the p-value of the aSPU test statistic by comparing the observed value 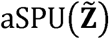 to the simulated values 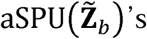 under the null, which gives us 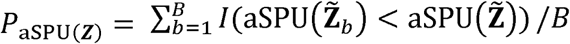. Literature has shown that by combining a variety of tests, the aSPU test can perform well in various scenarios, whereas conventional tests are usually not as robust (Pan and others, 2014; Kim and others, 2015).

### 2.4. General Procedure

In this section, we present the general procedure of applying our new approach to study the causal relationship between an exposure and multiple outcomes, as shown in Figure 2. We would like to point out that the second step is very crucial to MR analysis, since including SNPs that violate the IV assumptions may lead to problematic results. Nevertheless, for this manuscript, we emphasize on the fifth step, mainly comparing the multivariate approach with the univariate approach for modelling the relationship between the exposure and outcomes. We would also like to note that when conducting two-sample MR, it is more common to apply various MR techniques that use summary statistics (e.g. MR-Egger). We choose to focus on the two-stage MR method that is more comparable to our new multivariate approach, which requires individual level data. Another reason why we would like to focus on the two-stage methods is that two-stage regression allows correlated instrumental variables in the first stage model, whereas for summary-based MR methods, the chosen instrumental variables are required to be independent. In situations where there are not many available instrumental variables that are independent, it may be better to choose a looser correlation cutoff and apply the two-stage approach. We will discuss more on the strengths and drawbacks of our approach in the discussion section. Besides, it is worth mentioning that the two-stage methods can also be applied to the one-sample situation, where we have a single dataset containing information on G, X and Y. However, we would like to focus on the two-sample scenario, since literature has shown that only using one sample is less preferred, for it is more likely to result in biased estimates and inflated type I errors (Burgess and others, 2015; Xue and Pan, 2019). Some simulation results and discussion for the one-sample scenario are provided in Appendix A of the Supplementary materials.

**Fig. 2.**
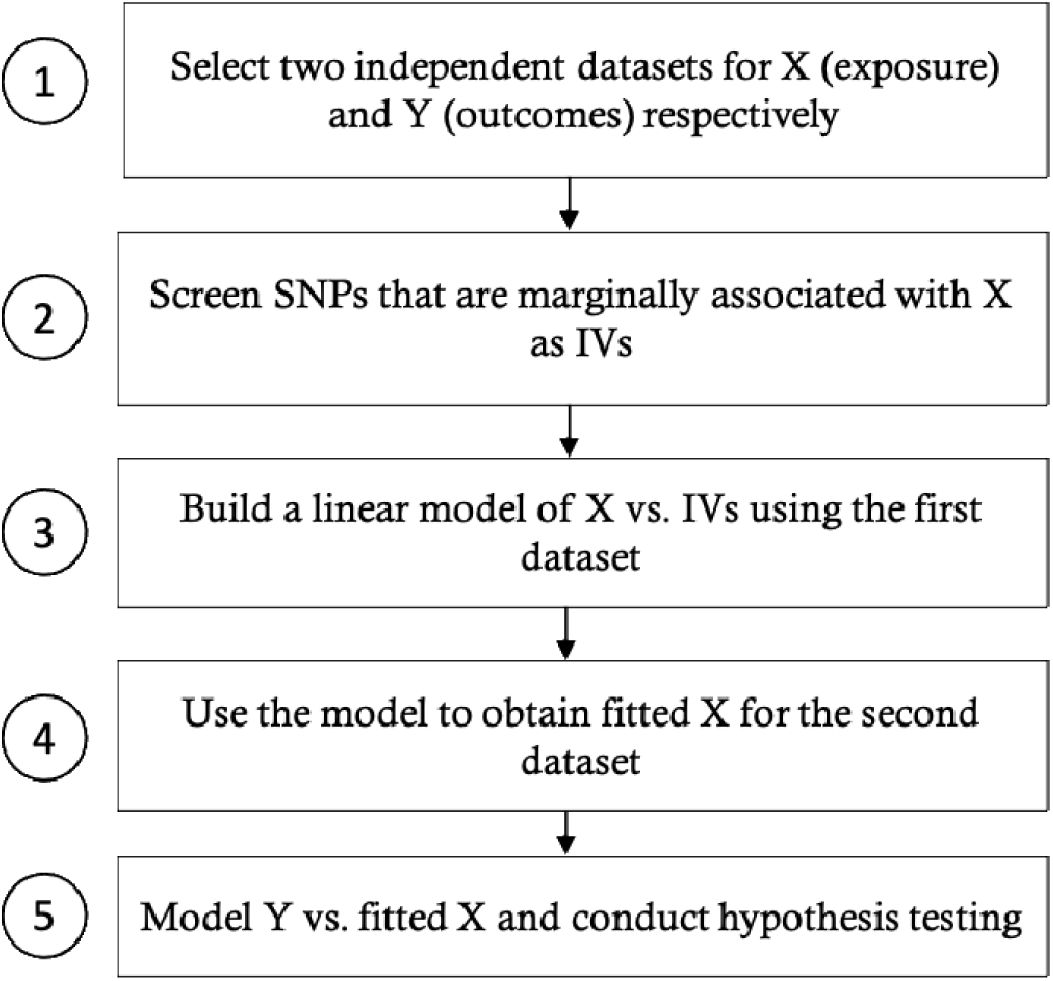
General procedure for applying MR to multiple outcomes using two samples with individual level data.

## 3. Results

### 3.1. Simulation studies

We conduct simulation studies to compare the performances of our new approach and the univariate two-stage MR method. Among the common variants that show some marginal association (p<1e-4) with baseline magnesium (MG) based on the CO.20 genotype data (Siu and others, 2013; Ringash and others, 2013; Shepshelovich and others, 2018), we randomly select 19 of them with weak or no correlations (pairwise correlations between -0.1 and 0.1). All of these SNPs have minor allele frequencies (MAFs) greater than 0.05, and 16 of them have MAFs greater than 0.1. In the simulations, we assume these SNPs are the true association SNPs, and simulate the genotypes for two independent samples, each with sample size 559, using multivariate binomial distributions and maintaining the MAF of each SNP. Since the correlations among the selected SNPs are very weak, for convenience, we simulate the SNPs as independent unless otherwise stated. Our experience shows that using weakly correlated SNPs or independent SNPs does not really affect the results. Suppose there are *q* = *q*_1_ + *q*_2_ outcomes. The first *q*_l_ outcomes are binary, while the other *q*_2_ outcomes are continuous.

We simulate U, X and Y using the following models:

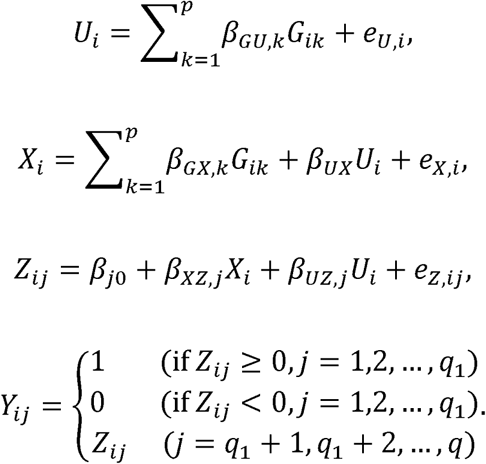

*G*_*ik*,_ *U*_*i*,_ *X*_*i*_, *Y*_*ij*_ *are* the *k*th genotype, confounder, exposure and *j*th outcome for subject *i.Z*_*ij*_ is the *j*th latent continuous outcome variable. For binary outcomes, *Y*_*ij*_ and *Z*_*ij*_ are connected by the probit link, where *Y*_*ij*_ follows a Bernoulli distribution with mean Ф(*Z*_*ij*_) (Ф is the cumulative density function of the standard normal distribution). For continuous outcomes, *Y*_*ij*_ = *Z*_*ij*_. *β*_*GU,k*_, *β*_*GX,k*_ are the effects of the *k*th SNP on U and X respectively. *β*_*XZ,j*_, and *β*_*UZ,j*_, are the effects of X and U on the *j*th latent variable respectively *β*_*UX*_ is the effect of U on X. *e*_*U,i*_, and *e*_*X,i*_,, are random errors that follow i.i.d. standard normal. ***e***_*z,i*_ *=* (*e*_*z,i*1_, …, *e*_*z,iq*_)′ follows an i.i.d. multivariate normal distribution with mean 0. The standard deviation of *e*_*z,ij*_, is σ_*j*_, and the correlation between *e*_*z,ij*_, and *e*_*z,il*_, is *ρ*_*jl*_ Denote the correlation matrix (*ρ*_*jl*_)_*q×q*_ by **Ω**. Also denote ***σ*** = (*σ*_1_, … *σ*_*q*_)′, ***β***_*GU*_ = (*β*_*GU*,1_, …, *β*_*GU,p*_)′, ***β***_*GX*_ = (*β*_*GX*,1_, …, *β*_*GX,p*_)′, ***β***_*XZ*_ = (*β*_*XZ*,1_, …, *β*_*XZ,p*_)′, ***β***_*UZ*_ = (*β*_*UZ*,1_, …, *β*_*UZ,p*_)′.

Following Slob and Burgess (2020), we generate *β*_*GX,k*_’s from a normal distribution with mean zero and standard deviation 0.15 first, and subsequently select those with *β*_*GX,k*_ > 0.08 to avoid weak IVs. Then we scale *β*_*GX,k*_ ’s by a constant so that the proportion of X explained by IVs is about 20%. For simplicity, we assume *β*_*GU,k*_ ’s are 0. *β*_*UX*_ and *β*_*UZ,j*_’s follow a standard uniform distribution.

At first, we consider 2 binary and 2 continuous outcomes. We set ***σ*** = (1,1,2,3)′ To make sure different binary outcomes have different event rates, we select ***β***_0_ = (−0.4, −0.8,0,0)^′^ For the continuous outcomes, *β*_*j*0_’s do not influence the results. In terms of the correlation structure, we let

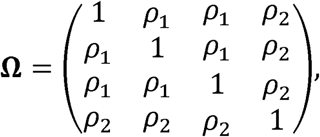

meaning that the first three outcomes have an exchangeable correlation structure with parameter *ρ*_l_, while the last outcome’s correlation with each of the first three outcomes is *ρ*_2_ After we simulate the two-sample data, we compare univariate and multivariate two-stage analyses in terms of type I errors and power. Next, we explore how univariate and multivariate methods perform when there are more mixed outcomes. We randomly select 7 outcomes (3 binary, 4 continuous) from the CO.20 data and use their correlation structure as our **Ω**. In this case, we select ***σ*** = (1,1,2,3,4,5)^′^ and ***β***_0_ = (−0.2, −0.5, −0.9,0,0,0,0)^′^.

As Figure 3 shows, both univariate and multivariate analyses are able to control type I errors when testing the overall hypothesis or each single outcome. In terms of power, we examine various scenarios, for which the different parameter settings are included in Table 1. Though we mainly focus on our proposed gradient descent algorithm for this manuscript, our experience shows that the results of gradient descent and Newton’s method are very consistent under most circumstances, as shown in Figure 4 for example.

**Table 1.**
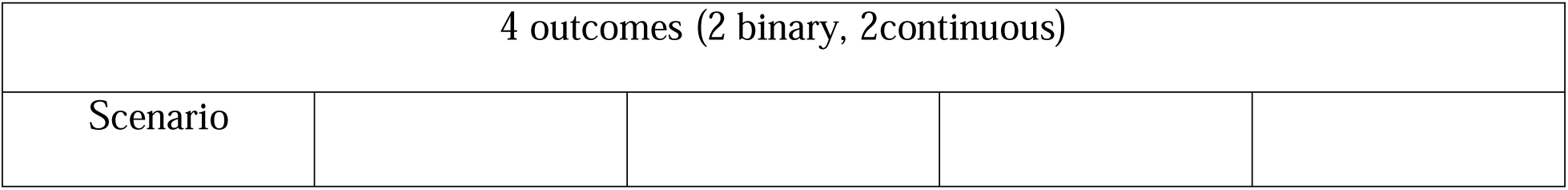
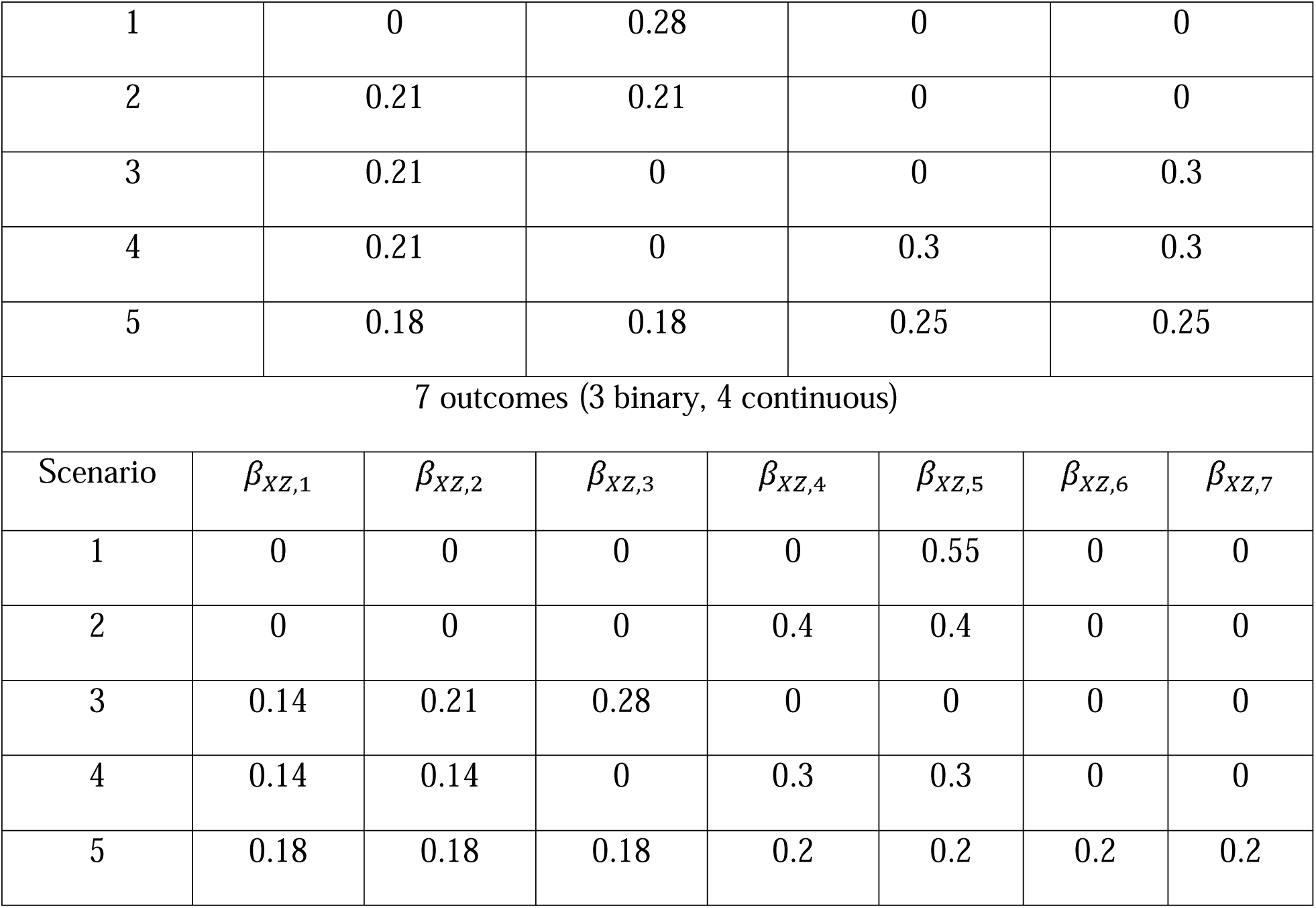
Different scenario settings used for power comparison.

| 4 outcomes (2 binary, 2continuous) |
| --- |
| Scenario |

|  |  |  |  |  |  |  |  |
| --- | --- | --- | --- | --- | --- | --- | --- |
| 1 | 0 | 0.28 | 0 | 0 |  |  |  |
| 2 | 0.21 | 0.21 | 0 | 0 |  |  |  |
| 3 | 0.21 | 0 | 0 | 0.3 |  |  |  |
| 4 | 0.21 | 0 | 0.3 | 0.3 |  |  |  |
| 5 | 0.18 | 0.18 | 0.25 | 0.25 |  |  |  |
| 7 outcomes (3 binary, 4 continuous) |  |  |  |  |  |  |  |
| Scenario | $\beta_{xz,1}$ | $\beta_{xz,2}$ | $\beta_{xz,3}$ | $\beta_{xz,4}$ | $\beta_{xz,5}$ | $\beta_{xz,6}$ | $\beta_{xz,7}$ |
| 1 | 0 | 0 | 0 | 0 | 0.55 | 0 | 0 |
| 2 | 0 | 0 | 0 | 0.4 | 0.4 | 0 | 0 |
| 3 | 0.14 | 0.21 | 0.28 | 0 | 0 | 0 | 0 |
| 4 | 0.14 | 0.14 | 0 | 0.3 | 0.3 | 0 | 0 |
| 5 | 0.18 | 0.18 | 0.18 | 0.2 | 0.2 | 0.2 | 0.2 |

**Fig. 3.**
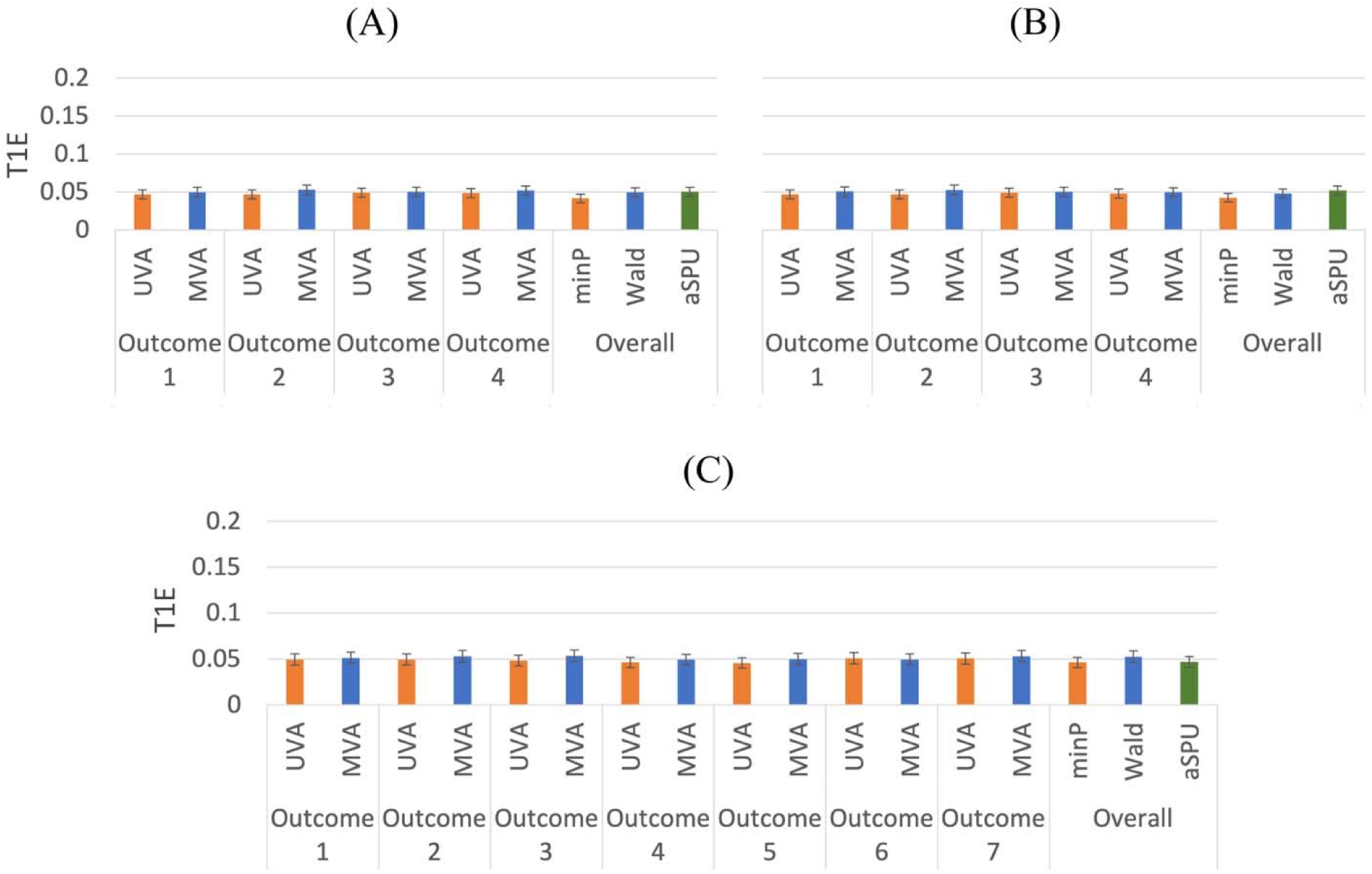
Type I error comparison of univariate and multivariate methods for mixed outcomes. 5000 replications. (A) 2 binary and 2 continuous outcomes. *ρ*_1_ = 0.3, *ρ*_2_ = 0.5. (B) 2 binary and 2 continuous outcomes. *ρ*_1_ = 0.3, *ρ*_2_ = −0.5. (C) 3 binary and 4 continuous outcomes. *β*_*XZ*,1_ = *β*_*XZ*,2_ = … =*β*_*XZ*,7_ = 0.

**Fig. 4.**
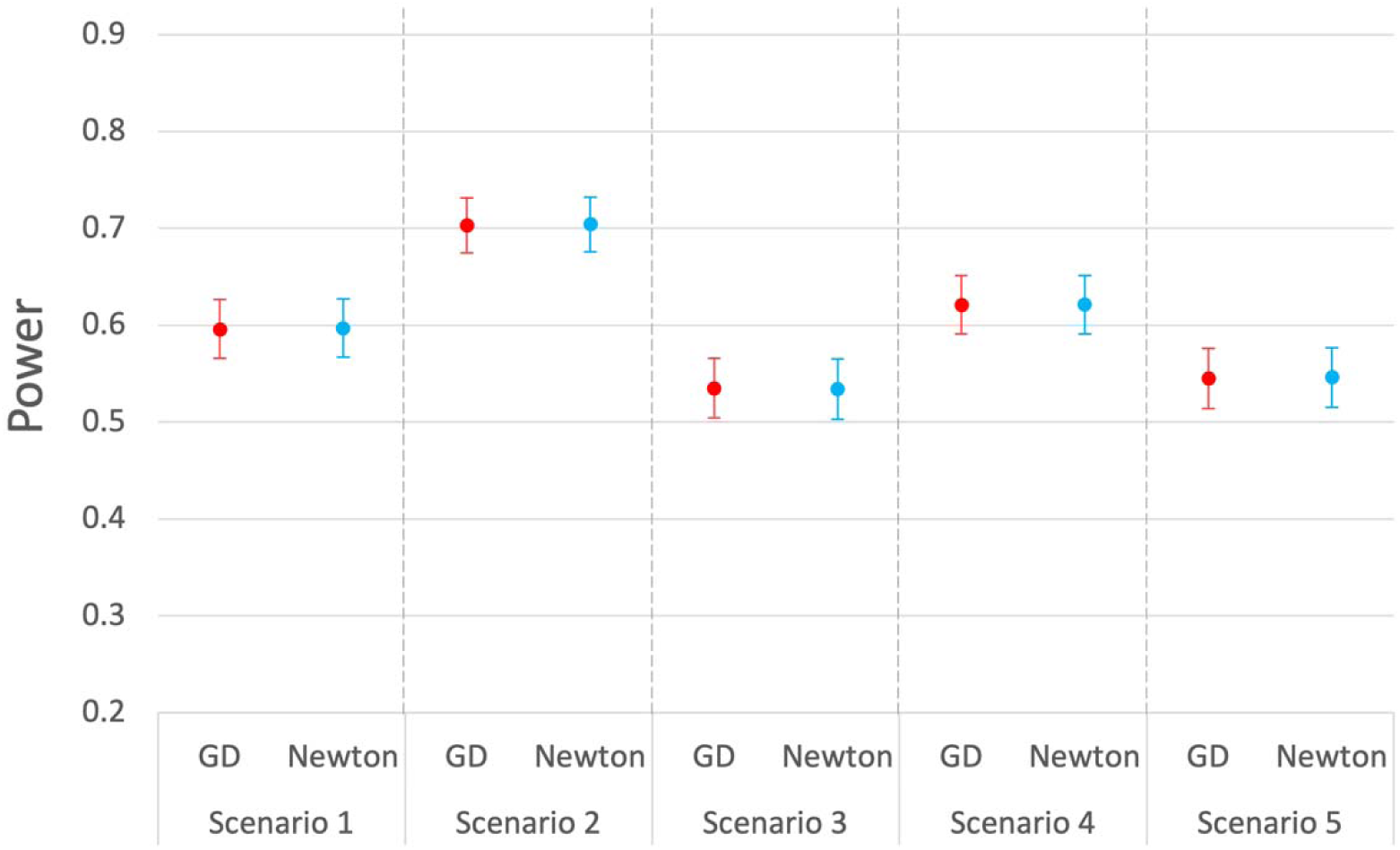
Power comparison of the Wald tests for mixed outcomes (2 binary, 2 continuous) using gradient descent and Newton’s method. 1000 replications.

Comparing the power performances of univariate and multivariate analysis with regard to the overall test (minP for univariate analysis; Wald, aSPU for multivariate analysis), we find that the Wald test usually has higher power than the minP test, as shown in Figure 5. The aSPU test has the highest power when the proportion of outcomes affected by the exposure is relatively high, though it may be disadvantaged when there is only one outcome affected by the exposure. Our finding is consistent with the literature about the aSPU test (Kim and others, 2015). The minP test is close to the SPU(∞) test except that it does not use the correlation information, and thus it works relatively well when the signal is sparse (e.g., the exposure only has an effect on one of the outcomes). The Wald test is similar to the SPU(2) test, which performs the best when the signal is not very sparse. This may partially explain why the overall tests using multivariate analysis tend to show a larger power improvement over the minP test when the exposure affects more than one outcome.

**Fig. 5.**
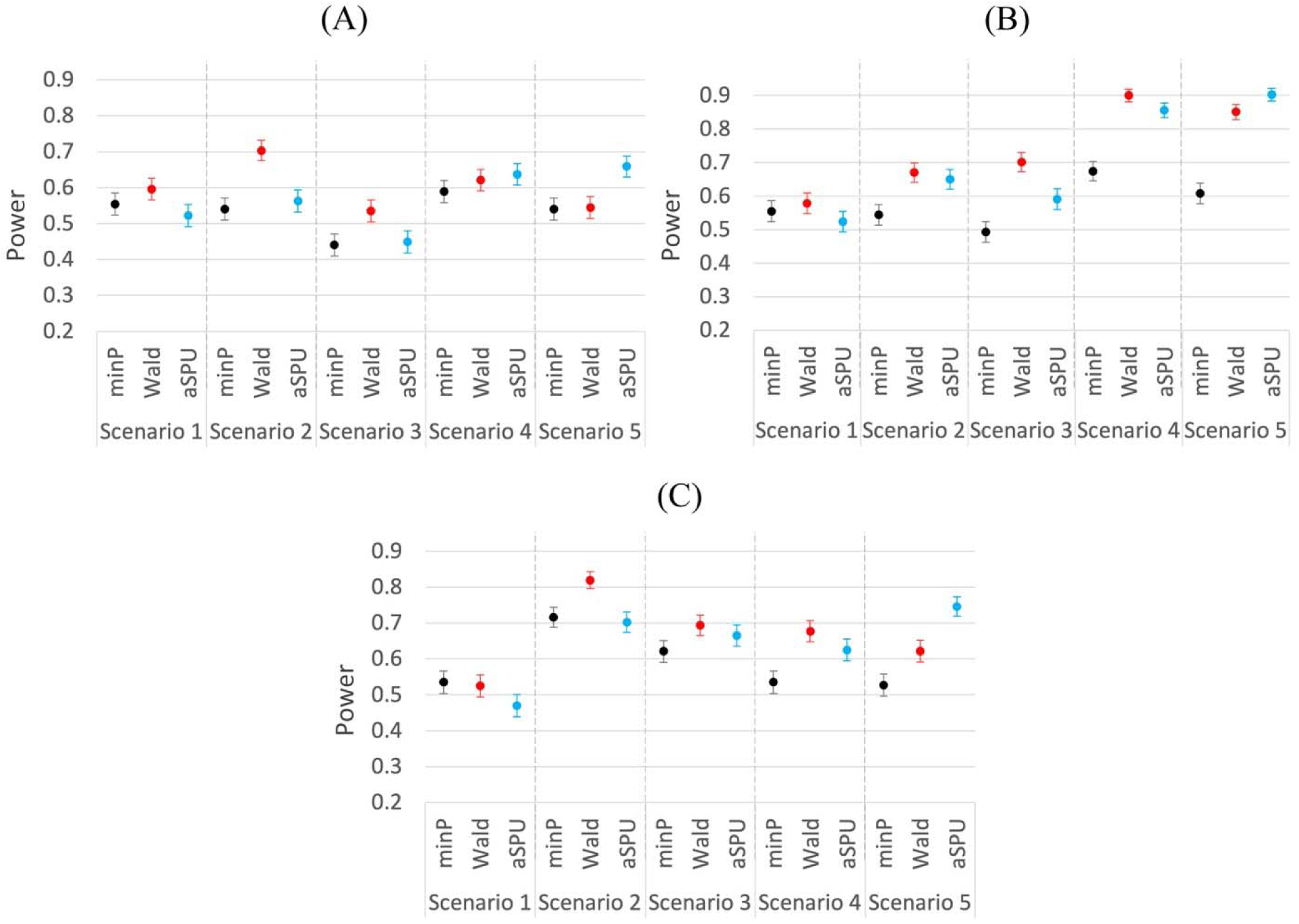
Power comparison of univariate and multivariate methods for mixed outcomes. 1000replications. (A) 2 binary and 2 continuous outcomes. *ρ*_1_ = 0.3, *ρ*_2_ = 0.5. (B) 2 binary and 2 continuous outcomes. *ρ*_1_ = 0.3, *ρ*_2_ = −0.5. (C) 3 binary and 4 continuous outcomes.

Note that for the above scenarios, we assume that the IVs are independent, which is usually the case for Mendelian randomization studies using summary statistics. However, for the two-stage methods, we can demonstrate that IVs with moderate correlations can also be used. We explore another situation with 4 mixed outcomes (2 binary, 2 continuous) and 10 correlated IVs. We select *ρ*_1_ = 0.3, *ρ*_2_ = 0.5, and assume the correlated IVs have an AR(0.4) correlation structure, meaning that the correlation between the *k*_l_ th SNP and the *k*_2_ th SNP is defined as 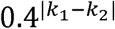. Each SNP’s MAF is randomly chosen using Unif(0.3, 0.5). The other settings are the same as before. According to Figure 6, when the IVs are moderately correlated, both univariate and multivariate two-stage methods are still able to control type I errors. We also observe similar power performances to those in the previous simulation settings. Again, the Wald and aSPU tests outperform the minP test when there is more than one outcome affected by the exposure, and the aSPU test has the most advantage when all outcomes are affected. More simulation results with modified settings are included in Appendix A of the Supplementary materials.

**Fig. 6.**
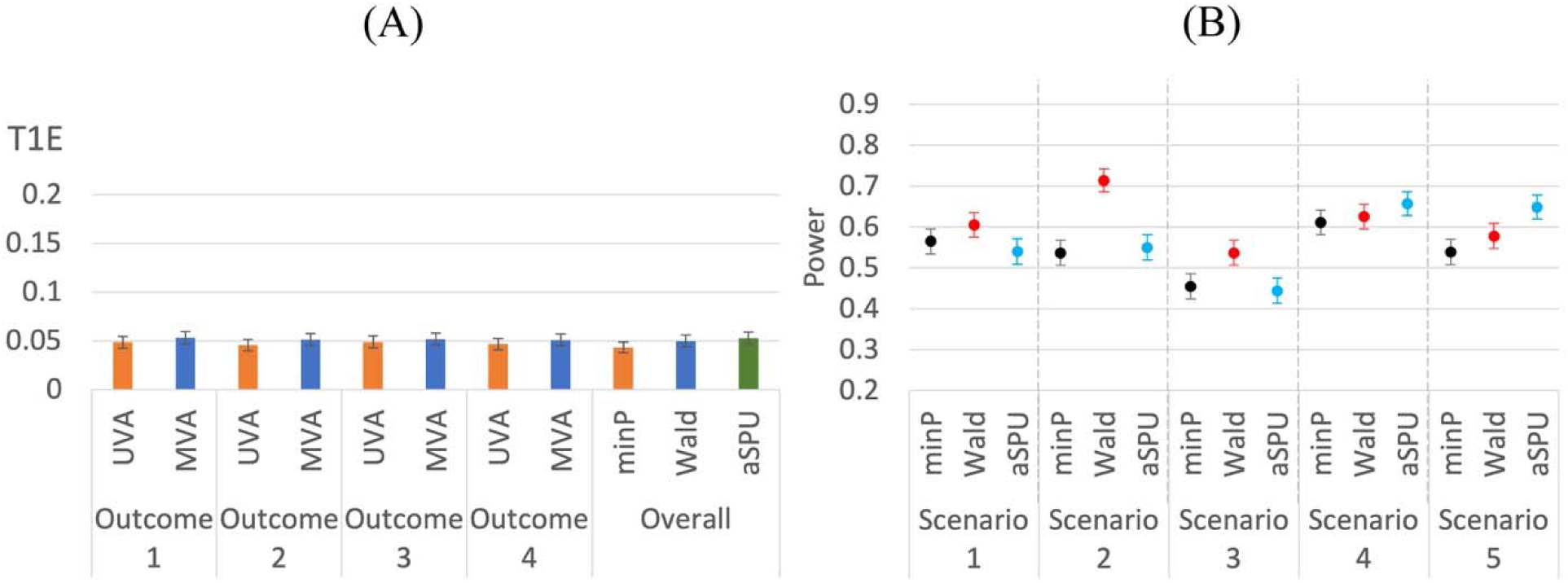
T1E and power comparison of univariate and multivariate methods for mixed outcomes using 10 correlated IVs.. (A) T1E, 5000 replications. (B) Power, 1000 replications.

### 3.2. Real Data Application

To further demonstrate the difference of univariate and multivariate analyses in practice, we look at the data from CO.20 trial as mentioned in the introduction, which randomly assigned into two treatment groups (cetuximab + B\ri; cetuximab + placebo), which, for convenience, we denote as Group 1 and Group 2 respectively. We choose baseline magnesium level (scaled by 10) as our exposure of interest X, since it is associated with certain genetic variants, and we are interested in examining whether this variable affects any of the following outcome variables: 3 binary toxicity variables (rash, nausea and vomiting) and 4 continuous lab variables (bilirubin [BIL], white blood count [WBC], alanine aminotransferase [ALT] and lactate dehydrogenase [LDH]). Each binary toxicity variable is coded as 0 and 1 (has not experienced or has experienced a toxicity event within 8 weeks), and each lab variable is defined as the worst (maximum) value within 8 weeks after allocation. We take log-transformation on BIL, ALT and LDH and remove 3 subjects with outlying values, defined as more than 4 standard deviations away from the sample mean value. The sample correlation matrix of the 7 outcomes is illustrated in Appendix B of the Supplementary materials, which shows that most outcomes are weakly or moderately correlated.

Using the data from Group 1, we obtain the marginal associations of SNPs with X (adjusted for age and gender) and select SNPs with marginal p-values smaller than 1e-4 and MAFs greater than 0.05. After pruning the SNPs to control pairwise correlations to be within -0.1 and 0.1, we end up with 6 SNPs as our IVs and build a first stage model of X vs. IVs. Next, we use the first stage model to obtain fitted X for Group 2, after which we conduct the second stage analysis, applying the univariate and multivariate modeling approaches to explore whether X has any effect on any of the mixed outcomes. Figure 7 shows a summary of our workflow.

**Fig. 7.**
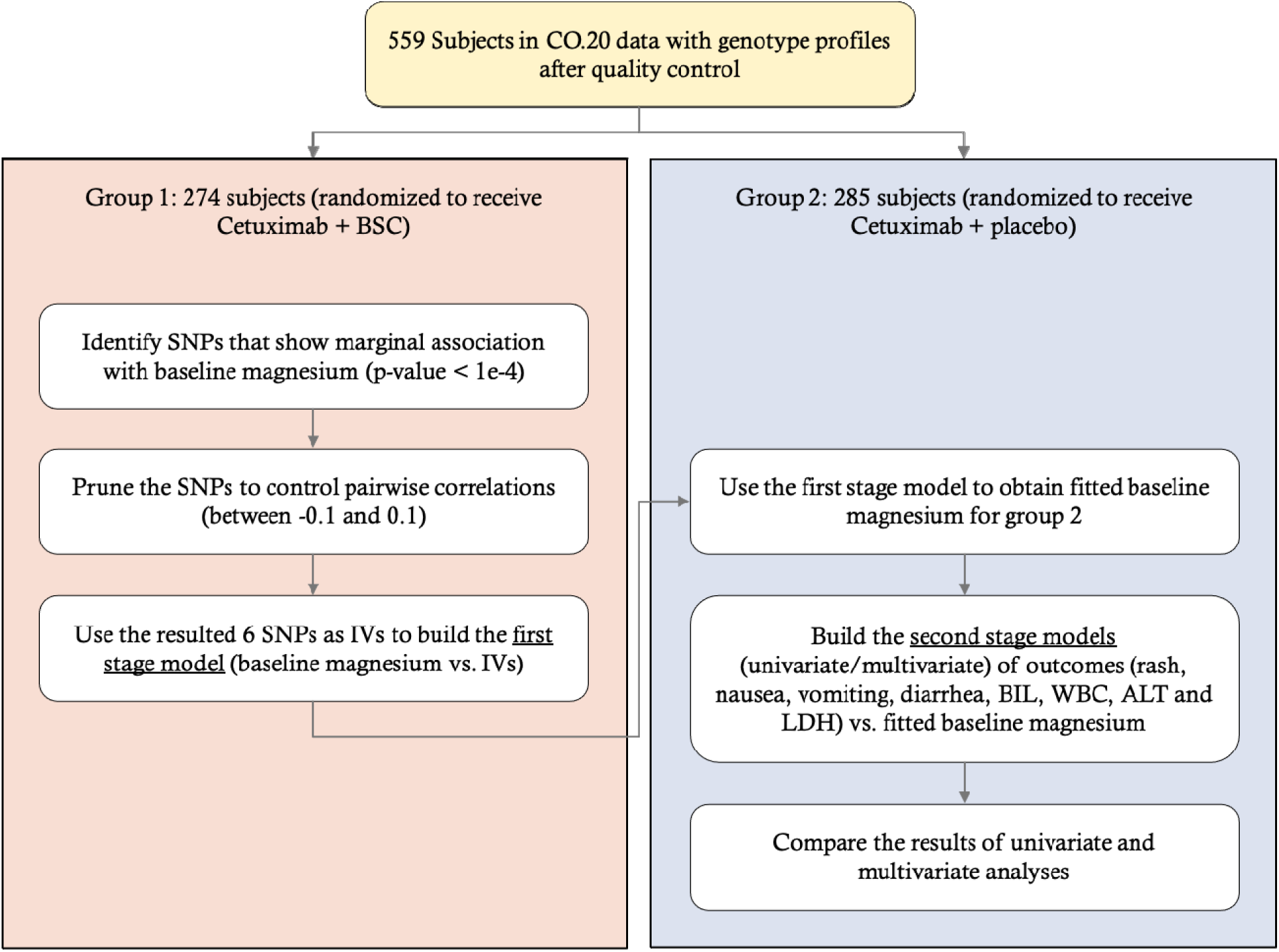
Workflow for the CO.20 study application.

The p-values for testing baseline magnesium’s effect on each of the outcomes based on univariate analysis are shown in Table 2. We also include the result of univariate analysis without MR, where each outcome is regressed on observed X instead of fitted X. According to the table, based on univariate MR analysis, BIL and ALT show some significance with p-values less than 0.05. Meanwhile, possibly as a result of unhandled confounders, the p-values based on univariate analysis without MR are usually quite different. This may partially demonstrate the effect and importance of considering instrumental variables for causal inference.

**Table 2.** P-values for testing baseline magnesium’s effect on each of the outcomes based on two-stage univariate MR analysis and univariate analysis without MR. Exposure: baseline MG.

|  | Rash | Nausea | Vomiting | BIL | WBC | ALT | LDH |
| --- | --- | --- | --- | --- | --- | --- | --- |
| MR | 0.078 | 0.172 | 0.210 | 0.010* | 0.289 | 0.033* | 0.238 |
| No MR | 0.855 | 0.013* | 0.786 | 0.071 | 0.511 | 0.019* | 0.477 |

Table 3 summarizes the results of univariate and multivariate analyses. For estimating and testing a single effect, univariate and multivariate analyses provide consistent results, though they are not exactly the same. However, the overall hypothesis test results are very different. The p-value of the minP test is not significant, but the p-values of the Wald and aSPU tests are. This agrees with our simulation results, demonstrating the potential power advantage of multivariate analysis.

**Table 3.** Comparison between univariate and multivariate methods for mixed outcomes (rash, nausea, vomiting, BIL, WBC, ALT and LDH). Exposure: baseline MG.

|  | UVA |  |  | MVA (MCLE) |  |  |
| --- | --- | --- | --- | --- | --- | --- |
|  | Effect size | SE | P-value | Effect size | SE | P-value |
| Rash | -0.266 | 0.151 | 0.078 | -0.267 | 0.154 | 0.083 |
| Nausea | -0.224 | 0.164 | 0.172 | -0.224 | 0.164 | 0.171 |
| Vomiting | 0.248 | 0.198 | 0.21 | 0.259 | 0.191 | 0.174 |
| ALT | -0.159 | 0.074 | 0.032* | -0.159 | 0.074 | 0.032* |
| BIL | -0.191 | 0.074 | 0.01* | -0.191 | 0.074 | 0.01* |
| LDH | -0.093 | 0.079 | 0.237 | -0.093 | 0.072 | 0.193 |
| WBC | -0.446 | 0.42 | 0.289 | -0.446 | 0.351 | 0.204 |
|  | minP |  |  | Wald |  | aSPU |
| Overall | 0.07 |  |  | 0.015* |  | 0.021* |

Thus, the proposed MRMO model is shown to identify the statistically significant associations between genetic predisposition of having lower magnesium levels to hepatotoxicities (elevated ALT levels and elevated bilirubin levels) in cetuximab-treated patients.

## 4. Discussion

We have presented a novel approach to conduct two-stage Mendelian randomization with multivariate analysis on mixed outcomes. As shown in our simulations, our innovative approach can increase the power of the overall test over the univariate approach in most scenarios, especially when more than one outcomes are affected by the exposure. The two different overall tests based on multivariate analysis, Wald and aSPU tests, have different performances in different scenarios. The aSPU test shows better power when the proportion of outcomes affected by the exposure is higher, while the Wald test performs the best when the exposure has causal effects on a small to medium proportion of the outcomes. We have also noticed that carrying out the tests using our proposed gradient descent algorithm tends to yield very similar results to those based on the Newton-Rapson method, which was used by Bai and others (2020). Nevertheless, given the possible drawbacks of the Newton-Rapson method in some other scenarios (Press and others, 1988; Strutz 2010; Pascanu and others, 2014), we recommend using the gradient descent approach by default, which may take more iterations to converge but can avoid some potential computation issues.

After applying the univariate and multivariate methods to the CO.20 data, we have found that the parameter estimations for single outcomes were close. However, the minP test for testing the overall hypothesis did not give a significant p-result, whereas both the Wald test and the aSPU test in our new method had significant results. The increased statistical power offered by multivariate analyses was therefore able to identify a previous latent relationship between predisposition to hypomagnesemia and hepatotoxicity from cetuximab-treated chemo-refractory colorectal cancer patients. This is relevant clinically because hypomagnesemia has been associated with improved outcomes in cetuximab-treated patients. Because we identified that predisposition to low magnesium may lead to increased liver toxicities, new avenues of research have been opened, where predicposition to low magnesium levels may lead either to increased levels of cetuximab (pharmacokinetic association) or increased efficacy of cetuximab on the metastatic cancer (pharmacodynamic association). Confounding this association is that knowledge that elevated bilirubin may be due either to drug therapy or bulky liver metastasis. Because we also found a separate relationship between predisposition to hypomagnesemia and elevated ALT, this result may suggest but does not prove that the relationship may be related more to drug toxicities than liver metastases, because ALT abnormalities are more commonly seen with drug toxicity than with liver metastases.

We would like to point out that our chosen screening criteria for the instrumental variables were relatively loose due to the limited sample size of the CO.20 data, which may raise concerns about the possible violation of instrumental variable assumptions. In order to alleviate this potential problem, one approach we can consider is to combine different datasets to get a larger sample size. This may require modifying our model framework to take the different structures of different datasets and potential correlations into account, which is a future direction worth exploring. Another possible approach is to use publicly available genome-wide association study (GWAS) results, which are usually based on sufficiently large samples, to select instrumental variables. However, it will not be feasible when there is no available GWAS results on the exposure of interest.

In the future, we may explore the possibility of developing multivariate methods that only requires summary statistics, as well as methods that can handle certain invalid instrumental variables. Besides, as we have demonstrated, our current approach can handle a mixture of binary and continuous outcomes. It can be directly applied to situations where all outcomes are binary, or all outcomes are continuous as well. We may also look at other types of outcomes (e.g., survival outcomes) and develop new methods that incorporate multivariate Mendelian randomization.

## 5. Software

R code and simulation data are available at https://github.com/yangq001/MRMO.

## Acknowledgments

The authors would like to acknowledge the clinical contributions of Lillian Siu.

