## Supplementary materials for "Two-Stage Multivariate Mendelian Randomization on Multiple Outcomes with Mixed Distributions"

### Appendix A, Additional simulation settings and results

We conduct additional simulations to study the performances of two-stage univariate and multivariate MR methods under various circumstances. For convenience, we refer to the setting with 4 outcomes and independent SNPs in the main article as setting 0. By default, we select $\rho_{1}=0.3, \rho_{2}=0.5$. For type I error evaluation, we use 5000 replications. For power evaluation, we use 1000 replications.

#### Setting 1: No correlation between outcomes

The only difference between setting 1 and setting 0 is that for setting 1, the 4 outcomes are independent, meaning that $\rho_{1}=\rho_{2}=0$. As shown in Figure S1, type I errors are controlled, and we observe power patterns similar to what we see under setting 0. Under this new setting, the minP test for univariate analysis performs the best in scenario 1, where only one outcome is affected by the exposure. One possible explanation is that when there is no correlation between the outcomes, multivariate analysis does not benefit from using the correlation information as much as it does when the outcomes are correlated. Instead, it may slightly lose power since it still needs to estimate the parameters related to correlations. Nevertheless, when there are multiple outcomes affected by the exposure, multivariate analysis still shows higher power than univariate analysis.

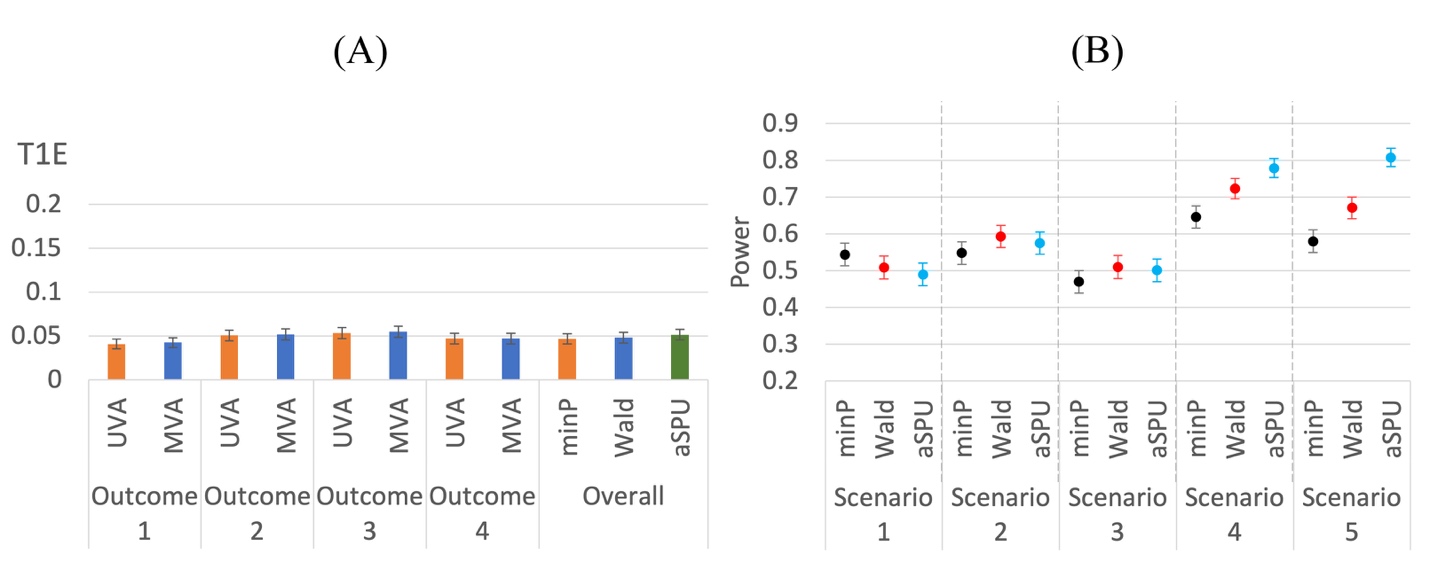

**Fig.S1.** T1E and power comparison of univariate and multivariate methods for mixed outcomes under setting 1.

#### Setting 2: Strong correlations between outcomes

The only difference between setting 2 and setting 0 is that for setting 2, $\rho_{1}=0.4,\rho_{2}=0.6$. As shown in Figure S2, both univariate and multivariate analyses are able to control type I errors. We observe similar power advantages of the multivariate analysis to those we demonstrate in setting 0. The extent of power increase can be larger when the correlations are higher, which is reasonable, since univariate analysis is suffering more by ignoring the correlation information.

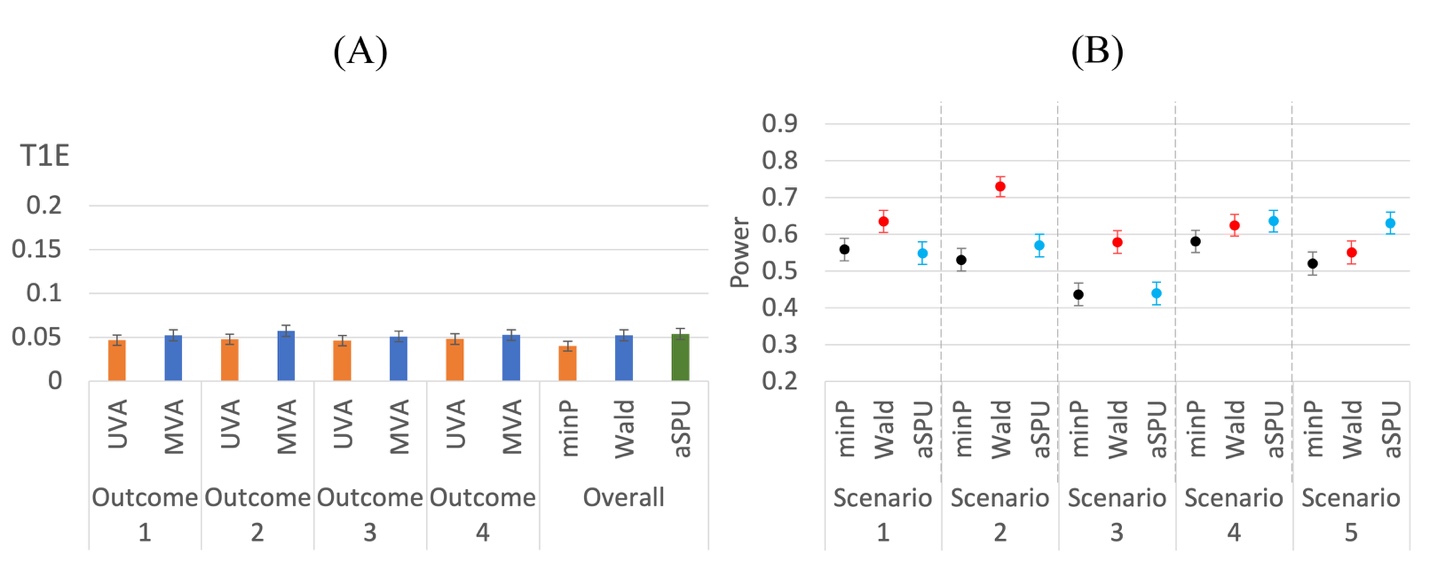

**Fig.S2.** T1E and power comparison of univariate and multivariate methods for mixed outcomes under setting 2.

#### Setting 3: One sample

The only difference between setting 3 and setting 0 is that for setting 3, we look at the one-sample setting, where a single sample of 559 subjects is used in both stages of the two-stage MR methods. As shown in Figure S3, type I errors are sometimes slightly inflated, which is consistent with the findings by Xue and Pan (2019). Hence, we highly recommend using the two-stage approaches with two samples if possible.

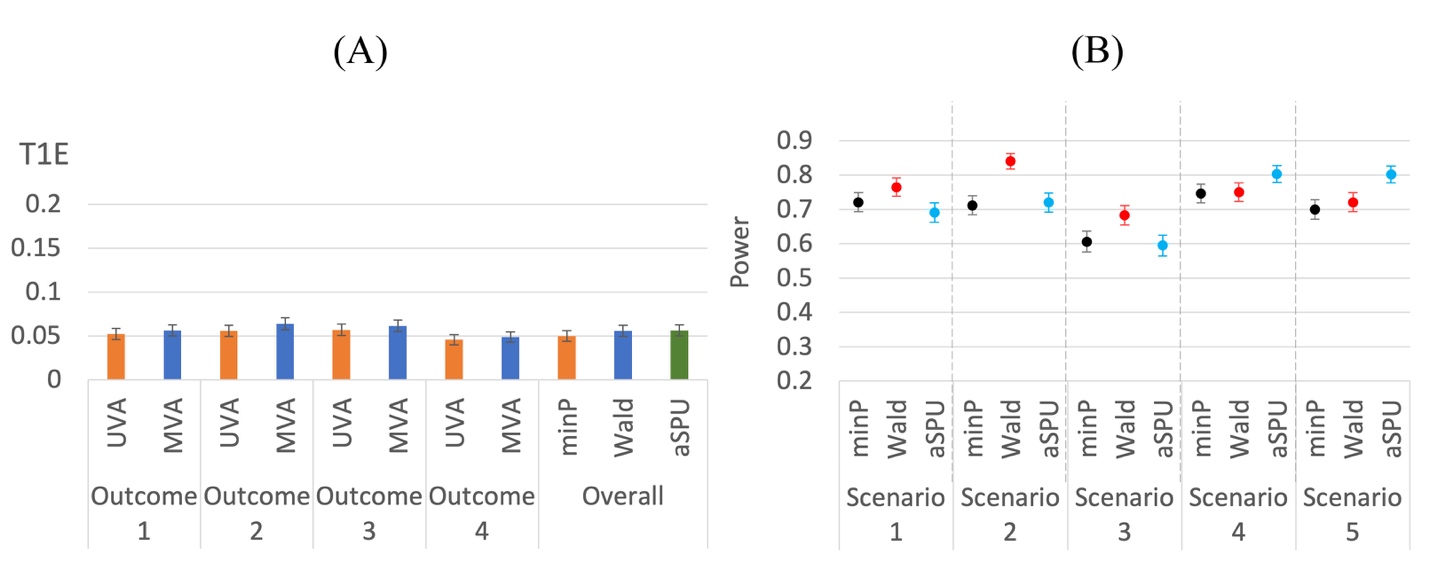

**Fig.S3.** T1E and power comparison of univariate and multivariate methods for mixed outcomes under setting 3.

### Appendix B, Correlation matrix of the seven chosen outcomes in the CO.20 data

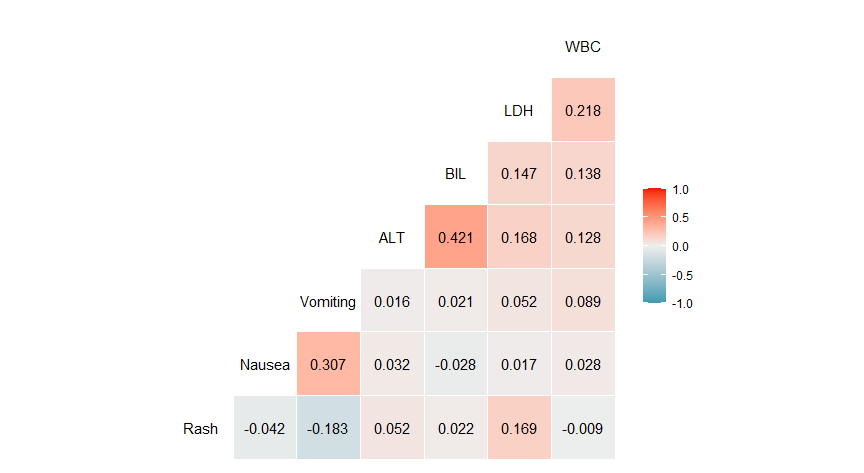

**Fig.S4.** Estimated correlations between different outcomes.
